# From external-input sensitivity to resident persistence: community assembly in a sink p-trap model

**DOI:** 10.64898/2026.05.13.724980

**Authors:** Qing Dai, Anthony A. Fodor, Ganchao Wei, Li Ma, Claudia K. Gunsch, Joshua A. Granek

## Abstract

Microbial habitats that receive repeated external input may not remain shaped by that input forever if local retention allows resident communities to build up over time. Here, we used a controlled bench-scale sink p-trap system to examine how community assembly unfolded during initial establishment in new, bleach-treated p-traps. Two p-traps received repeated handwashing-water input, while one received tap water as baseline. The treated p-traps, but not the control, showed clear successional change toward later resident-like states. Nested-model comparisons further showed that recent external input had its greatest influence early in succession, but the p-trap’s own prior state remained the stronger predictor throughout. Final-day post-flush trajectories indicated short-term displacement from pre-flush positions, with later time points tending to move back toward late-stage resident centroids. Together, these results show that repeated inoculation does not necessarily keep communities under continued outside influence. Instead, retentive microbial habitats can shift over time from early sensitivity to external input toward persistence shaped more by local history.

## Introduction

Community assembly after disturbance depends not only on which taxa arrive, but also on when they arrive and how strongly later community states remain shaped by assembly history^1^. Priority effects arise because the order and timing of immigration can alter the establishment success of later arrivals, making historical contingency a central feature of community assembly rather than a minor deviation from it^1,2^ . In microbial systems, succession is therefore not simply taxonomic replacement through time, but can reflect a shifting balance among dispersal, stochastic establishment, ecological selection, and legacy effects from prior community states^3^. Community development after disturbance may accordingly transition from relatively stochastic early establishment toward stronger deterministic or history-dependent structuring as succession proceeds^3^.

The distinction between input-sensitive assembly and history-dependent persistence has broader relevance for understanding how microbial communities respond to perturbation across engineered, host-associated and natural ecosystems. In the built environment, microbial communities are repeatedly perturbed by human use, cleaning, water flow, design, material properties, or other interventions, yet many built habitats support persistent microbial populations^4–7^. Understanding when these communities remain sensitive to incoming microorganisms and when they become increasingly constrained by local history is important for efforts to establish beneficial microbiomes in new built environments, shift undesirable communities toward healthier states, and increase resistance to colonization by opportunistic pathogens. Related questions arise in host-associated microbiomes, including the gut, where early colonization and priority effects can have long-term consequences for later community states, though these systems are additionally constrained by host physiology, immunity, diet, and other host-mediated selective pressures^8–10^. In natural ecosystems, analogous issues arise when communities respond to invasive species, disturbance, or restoration, where outcomes may depend on both the new arrivals and the historical state of the resident community.

Habitats that receive repeated external input while also permitting local retention and persistence may be especially informative for studying this transition. In such systems, present-day composition need not simply mirror the most recent input, because resident communities can increasingly constrain which later arrivals establish and persist^2,3^ . Sink p-traps provide a tractable habitat in which to examine this transition because they are repeatedly exposed to incoming water during use, can support persistent microbial populations, and often resist sustained decontamination, consistent with the possibility that local habitat structure and resident communities increasingly shape later states^11,12^ . Related work on plumbing-associated biofilms further shows that early colonization is dynamic and includes identifiable pioneer taxa, indicating that initial establishment and later community development can represent ecologically distinct phases^13^.

Longitudinal sampling of sink p-traps has shown that these communities develop through time and can re-establish after bleach treatment, consistent with temporal assembly and resilience rather than passive replacement alone^14^. However, most work on sink-associated systems has been framed in terms of pathogen reservoirs, recolonization, or intervention outcomes, leaving the broader assembly question less well resolved^11,14^ . A key unresolved question is whether communities continue to track recent external input or instead become increasingly shaped by within-habitat history as succession proceeds. Here, we used a controlled bench-scale p-trap system to examine microbial community assembly. By comparing p-traps receiving repeated handwashing-water input with a tap-water control, we asked whether p-trap communities follow a reproducible successional trajectory and whether maturation involves a shift from early external-input sensitivity to later history-dependent resident persistence. We then used overlapping shotgun metagenomic profiling to test whether the same dominant successional pattern was recoverable across sequencing modalities.

## Results

### Experiment Overview

To characterize the microbial colonization and community dynamics of new sink p-traps we tracked three new, bleach-treated bench-scale p-traps for 95 days: two treatment p-traps that repeatedly received handwashing water and one control p-trap that received tap water from the same faucet. This design allowed us to compare replicated treated p-traps against a shared-source control while following community development from early establishment to later maturation (Fig.1). Over the course of the experiment, p-trap water was sampled immediately before flushing, and samples of the incoming handwashing water and tap water were collected in parallel. On the final day, we added a higher-resolution time course by sampling immediately before flushing and again at 0, 2, 6, and 12 hours after flushing, followed by biofilm swabbing from the inner p-trap surface, to assess short-term disturbance and recovery.

**Figure 1.**
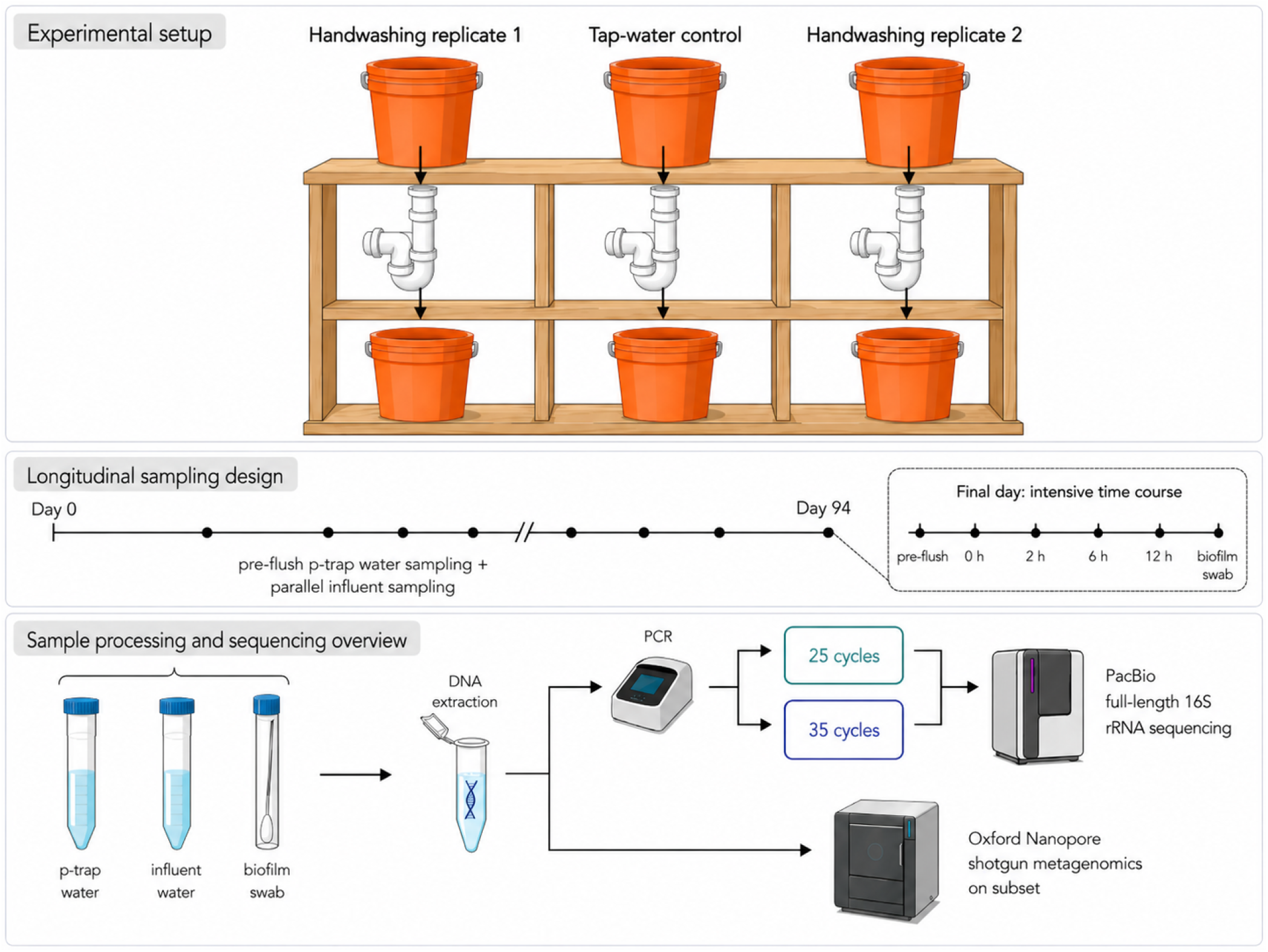
Experiment overview.

All samples were analyzed by PacBio full-length 16S rRNA amplicon sequencing. Biomass varied substantially and was particularly low for samples collected earlier in the experiment and shortly after flushing S5. For the lowest biomass samples DNA was undetectable after DNA extraction; for these samples a 35-cycle PCR amplification was used for library preparation. 25-cycle PCR amplification was used for the samples with detectable DNA. 42 of the samples with detectable DNA were included in both the 25-cycle and the 35-cycle amplification to allow comparison in order to characterize the effect of PCR bias under high levels of amplification. A subset of samples was also shotgun sequenced using Oxford Nanopore, allowing more detailed examination of microbial community structure.

Our analysis first examines the effect of high amplification (35-cycles) on observed community structure and whether the main ecological patterns were preserved between 25-cycle and the 35-cycle amplification. We then examined the successional trajectory of the p-trap communities, and finally tested whether maturation involved a shift from stronger early influence of recent external input toward later history-dependent resident persistence.

### Temporal community structure was reproducible across PCR-cycle conditions

Different library preparation conditions were used for the extremely low biomass samples (primarily early-stage samples) than for the moderate biomass samples (primarily mid- and late-stage samples). Library preparation for extremely low biomass samples, those with so little DNA that it was not detectable by Qubit fluorometry, used a 35-cycle PCR amplification, while a 25-cycle PCR amplification was used for moderate biomass samples. A subset of the moderate biomass samples (42 samples) were included in both the 25-cycle and 35-cycle library preparation batches.

We used this to subset of 42 sample subset to characterize the effect of additional PCR cycles, with the expectation that the 35-cycle library preparation would result in more PCR bias. We first asked whether the major ecological structure of the dataset was preserved. In the global PCA, samples clustered primarily by sample type and temporal position rather than by PCR-cycle batch, indicating that the dominant structure was maintained between amplification conditions (Fig. S1). Handwashing-water samples remained clearly separated from tap-water samples and p-trap water in both 25-cycle and 35-cycle datasets, while p-trap samples occupied a shared region of ordination space and spanned the main temporal gradient rather than forming amplification-specific clusters. Matched 25-cycle and 35-cycle samples (connected by lines in Fig. S1) were visibly displaced but remained embedded within the same higher-order ordination structure, indicating that PCR-cycle perturbed fine-scale positioning without losing the dominant high-dimensional structure associated with ecology.

Agreement between the datasets was even more clear when evaluating ASV trends over time. Among the 200 most abundant ASVs, directional temporal associations were broadly concordant between the 25-cycle and 35-cycle datasets, with many ASVs sharing temporal trends that are in the same direction (increasing or decreasing over time) between the datasets (Fig. 2). Thus, although amplification condition influenced fine-scale abundance structure, it did not erase the dominant temporal responses of abundant taxa.

**Figure 2.**
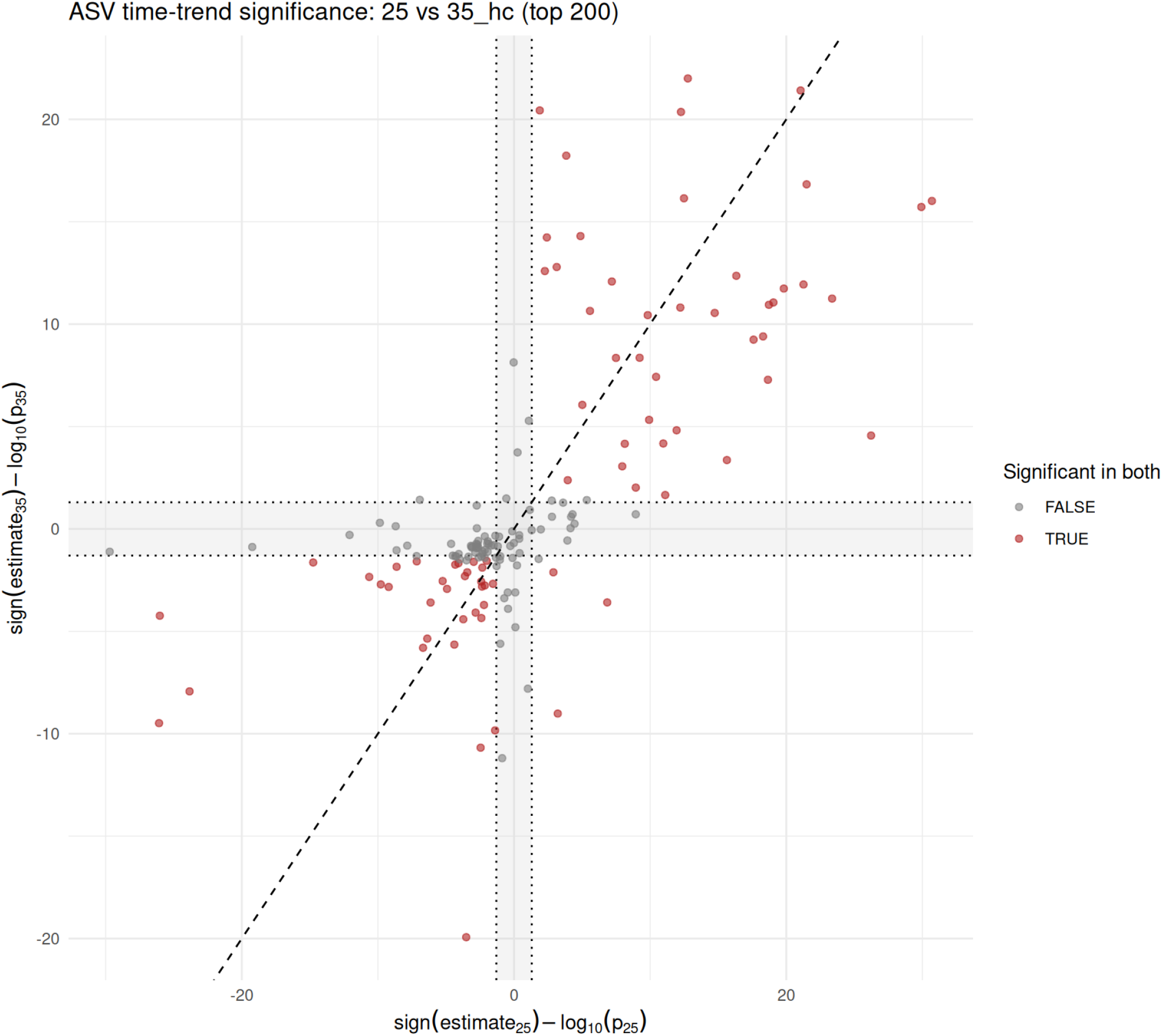
Concordance of ASV temporal associations across 25-cycle and 35-cycle datasets. Positive values of signed time-trend score indicate taxa that are increasing over time and negative values indicate taxa that are decreasing over time. Using mixed linear regression to test time trends of ASVs. Grey area are non-significant area.

Together, these analyses indicate that the differences between 25-cycle and 35-cycle amplifications affected sample composition less strongly than the major ecological signal, and that the dominant phenotype-level structure and temporal trends agreed between batches. For simplicity, the rest of our analysis focuses on the full 25-cycle dataset (not just the 42 sample subset shared between the 25-cycle and 35-cycle datasets). Results from the 35-cycle dataset are included as Supplementary Figures.

### P-trap communities changed along a shared dominant successional axis

We next examined the trajectory of the microbial community over time as microbes colonized the aseptic p-traps. PCA plots show a clear directional trajectory which appear to be correlated with sampling date, with samples from progressive weeks increasing along both PC1 and PC2 (Figs. 3 and S4). This pattern suggests succession from an early stage to late stage community. In the 25-cycle dataset, this directional trajectory is particularly strong in the two handwashing p-traps and less so for the tap-water p-trap. The time correlation is even more clear when we directly visualize principal component as a function of sample collection date and fit a generalized additive models (GAMs) (Fig. 4).

**Figure 3.**
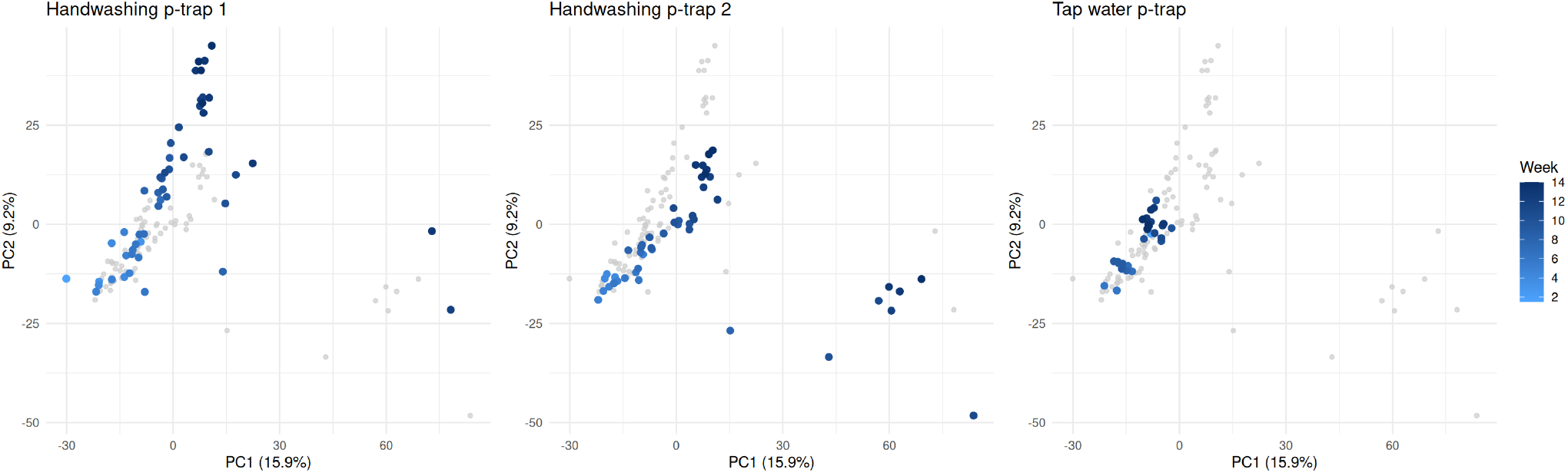
PCA trajectories show reproducible temporal progression in handwashing p-trap communities.

**Figure 4.**
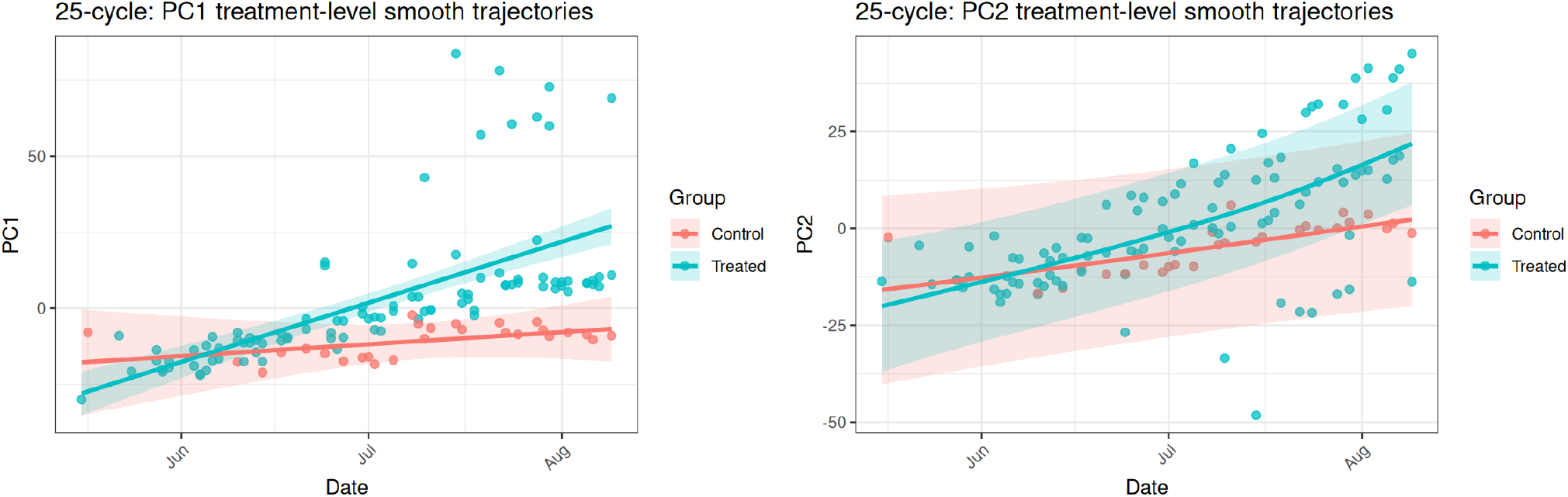
Treatment-level temporal trajectories along the major ordination axes in 25 cycle dataset. Points show sample scores, and lines with shaded bands show GAM-fitted trajectories ± 95% confidence intervals for treated and control p-traps.

The 35-cycle dataset recovered the same overall pattern despite a different axis orientation (Fig. S4B): both handwashing p-traps again followed strong, concordant displacement, whereas the tap-water followed a distinct and weaker trajectory.

### Community dynamics shifted from external-input sensitivity to history-dependent persistence

Having established that p-trap communities followed a similar successional trajectory, we next asked what best explained these changes at different stages of the experiment. To test this, we compared how well each sample could be explained by two possible sources of information: the most recent incoming water and the community already present in the same p-trap at the previous sampling time. We first fit a background-only model using blank samples, then asked how much the model improved when each source of information was added. Because mid-phase samples likely represented a transitional period, we focused our main comparison on early-versus late-phase samples. According to Fig. 5, adding information from the most recent incoming water produced a significantly larger improvement in model fit for early-phase samples than for late-phase samples, suggesting that newly forming p-trap communities were more sensitive to microbes entering with the water. By late succession, the same incoming-water information added much less explanatory power, indicating that each new input event left a smaller imprint on community composition. In contrast, information from the previous sample in the same p-trap remained strongly informative across phases and showed no significant early-to-late change. Together, these results suggest that p-trap communities shifted from an early stage in which incoming microbes had a stronger influence to a later stage in which community composition was more consistently tied to the community already present within each p-trap. This phase-dependent contrast was robust to alternative definitions of recent external input (Fig. S9).

**Figure 5.**
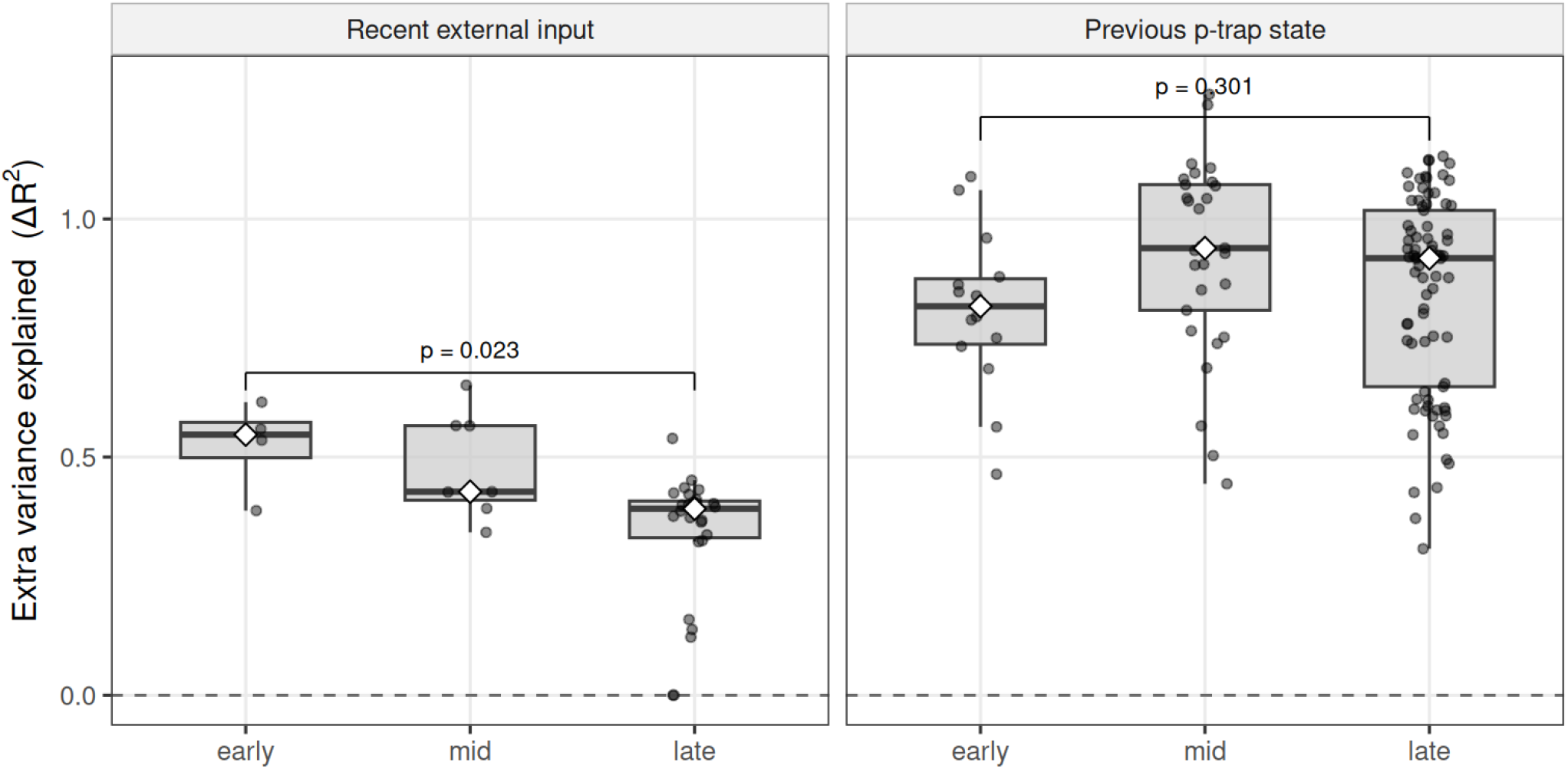
Community assembly shifted from external-input sensitivity to history-dependent persistence. Recent external input contributed most strongly during early assembly, whereas lagged within-p-trap state remained informative across phases and became dominant in later communities.

A short-timescale perturbation analysis supported the same interpretation. Final-day post-flush samples projected into the fixed 25-cycle PCA space showed transient displacement from their immediate pre-flush positions, but remained near, or moved back toward, the late-stage resident centroid over the subsequent hours (Fig. S11 and Fig. S12).Although post-flush movement varied among p-traps and was not uniformly monotonic, the overall pattern was consistent with short-term displacement around an already established late-stage resident community. Thus, the final-day flush appeared to perturb the community locally, while preserving the broader late-stage structure inferred from the time-series trajectory.

Together, these results indicate that succession in this p-trap habitat involved not only directional temporal change, but a shift in ecological control from externally sensitive assembly toward resident-governed persistence.

### Taxon-level attribution separated early colonizers from late persistent residents

We then asked whether this community-level shift was reflected in taxon-level patterns associated with early colonization versus late resident persistence. Taxon-level attribution analysis identified phase-dependent rankings of taxa who were better supported by recent external input during early succession or by the prior p-trap community state during late succession, consistent with a shift in the explanatory structure of taxon dynamics as p-trap communities matured.

In the 25-cycle dataset, taxa ranked highly in the early phase were dominated by organisms whose abundances were better explained when recent external input was added. The strongest representatives of this early colonization-associated cohort included the human-associated taxa *Staphylococcus epidermidis, Cutibacterium acnes*, and *Staphylococcus hominis*, together with taxa frequently detected in water-associated or low-biomass built-environment microbiomes, including *Methylobacterium* spp., *Sphingobium yanoikuyae*, and *Mycobacterium phocaicum* (Fig. 6A). These results indicate that, during early succession, recent input explained additional taxon-level variation beyond background signal, consistent with a community state that remained strongly coupled to external seeding.

**Figure 6.**
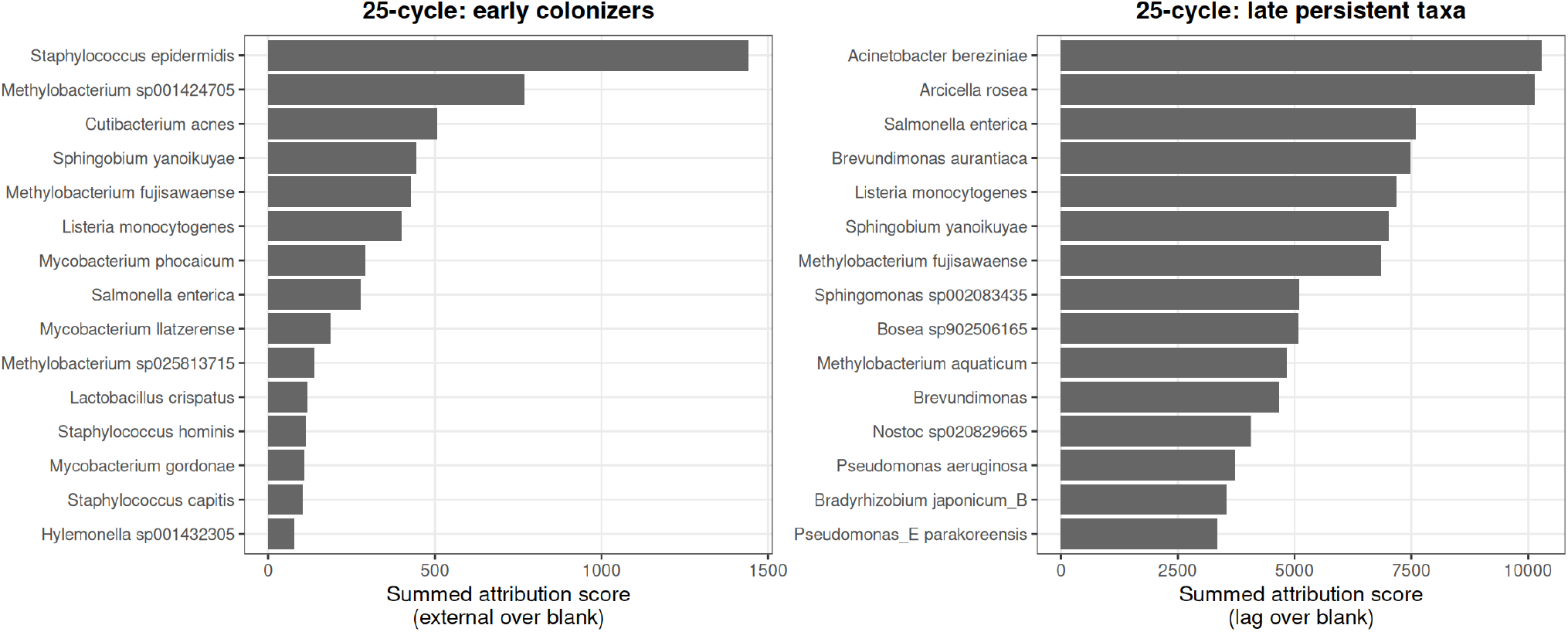
Taxon-level attribution identifies early colonizers and late persistent taxa in handwashing p-traps (25-cycle dataset). Left: Representative early colonizers in the handwashing p-traps. Right: Representative late persistent taxa in the handwashing p-traps.

By contrast, the highest-ranking late persistent taxa included *Acinetobacter bereziniae, Arcicella rosea, Salmonella enterica, Brevundimonas aurantiaca, Listeria monocytogenes, Sphingobium yanoikuyae*, and *Methylobacterium fujisawaense*(Fig. 6B).The partial overlap between early and late rankings indicates that the inferred transition was not simply a replacement of one taxonomic cohort by another, but a shift in the explanatory structure of taxon dynamics: recent input contributed more strongly during early succession, whereas late-stage abundances increasingly reflected the recent history of the p-trap community itself.

The 35-cycle dataset showed a similar phase-dependent attribution pattern, although the exact taxonomic rankings differed from the 25-cycle analysis (Fig. S10). In the early phase, taxa with the strongest external-over-blank attribution included human- or handwashing-associated organisms such as *Staphylococcus epidermidis, Enterococcus faecalis*, and *Cutibacterium acnes*, together with taxa frequently detected in water-associated or low-biomass built-environment microbiomes, including *Mycobacterium* spp., textitMethylobacterium spp., and *Novosphingobium* spp. These results indicate that, in the 35-cycle dataset, early taxon-level variation was again strongly supported by recent external input beyond the blank-only baseline.

In late stage, the highest-ranking history-associated taxa included *Acidovorax temperans, Acidovorax soli_A, Methylophilus* sp030696695, *Acidovorax* sp003058055, *Acinetobacter bereziniae, Afipia broomeae*, and *Arcicella rosea*. As in the 25-cycle dataset, some taxa appeared in both early and late rankings, indicating that the transition was not simply a complete replacement of one taxonomic group by another. Rather, the dominant explanatory signal shifted from recent external input during early succession to history dependence.

Taken together, taxon-level attribution across the 25-cycle and 35-cycle datasets supported a consistent ecological interpretation. Early communities contained taxa whose abundances were better explained when recent external input was included, consistent with stronger coupling to ongoing seeding during early assembly. By contrast, late-phase rankings emphasized taxa whose abundances were better explained by prior within-p-trap community composition, consistent with increasing historical dependence as resident communities became established. Although the specific taxa and their rankings differed between PCR-cycle conditions, both datasets converged on the same higher-order pattern: community maturation was accompanied by a shift in explanatory structure, from stronger external-input support early in succession toward stronger historical support in the late phase.

### Shotgun metagenomics independently recovered the dominant temporal pattern

To assess whether the temporal structure inferred from full-length 16S profiles was robust to sequencing approach, we generated long-read shotgun metagenomic data for an overlapping weekly subset of samples from handwashing p-traps. This analysis was intended as an orthogonal validation of the dominant temporal pattern identified in the 16S time series.

In both the full-length 16S and shotgun metagenomic datasets, the largest axis of compositional variation was organized by sampling time. Specifically, PC1 captured a broadly concordant temporal gradient across the overlapping weekly samples, with later samples separating from earlier samples along the dominant ordination axis in both p-traps and both sequencing modalities (Fig. 7). Thus, the primary structure of the data was not platform-specific: whether communities were profiled by full-length 16S amplicon sequencing or by shotgun metagenomics, the dominant compositional gradient corresponded to temporal progression.

**Figure 7.**
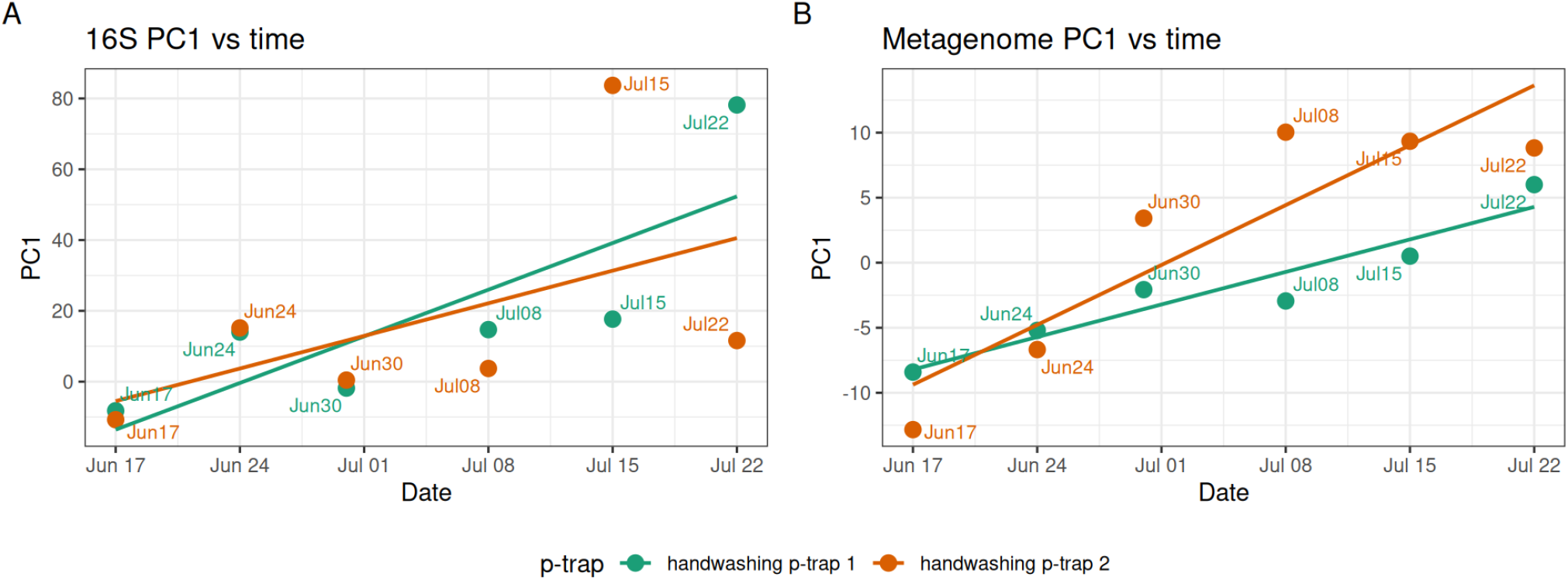
The dominant compositional gradient was organized by time in both amplicon and metagenomic profiles. (A) PC1 versus time in the full-length 16S dataset. (B) PC1 versus time in the shotgun metagenomic dataset.

Cross-platform agreement was also evident at the genus level. Among genera shared between the 16S and shotgun metagenomic datasets, genus-level associations with the platform-specific PC1 axes were positively correlated across platforms, and 75.0% of shared genera showed concordant effect direction (Fig. S8). Thus, the agreement between sequencing approaches was not limited to the ordering of samples along PC1; the genera contributing to the dominant temporal gradient were also broadly consistent between the two datasets.

This genus-level structure also showed agreement with the phase-specific attribution results. Several genera prioritized in the early external-input-associated attribution analysis, including *Staphylococcus, Cutibacterium*, and *Methylobacterium*, were negatively associated with PC1 in both the 16S and metage-nomic profiles, placing them on the early side of the temporal gradient. Conversely, several genera overlapping with the late history-associated attribution results, including *Bosea, Brevundimonas, Bradyrhi-zobium, Phenylobacterium*, and *Afipia*, were positively associated with PC1 in both datasets. Although genus-level PC1 associations are not expected to match species-level attribution rankings exactly, this concordance indicates that both sequencing approaches recovered similar taxonomic structure at the early and late ends of the dominant temporal axis.

Together, these results show that the dominant temporal structure inferred from full-length 16S profiles was also recovered by an independent long-read shotgun metagenomic readout. The agreement at both the sample level and genus level supports the interpretation that the leading axis of community variation reflected a reproducible successional gradient.

## Discussion

Our results support a broad ecological principle: in microbial habitats that receive repeated inputs while retaining established residents, community change can shift from input-driven assembly toward increasing dependence on resident community history. In the p-trap system studied here, communities followed directional successional trajectories toward later resident-like states, and this progression was accompanied by a change in the dominant ecological control on assembly. Early in succession, p-trap communities were more strongly associated with recent external input, whereas later communities became increasingly predictable from previous community state. The central transition in this system was therefore a shift in assembly logic from externally sensitive establishment toward increasingly history-dependent persistence.

This interpretation places sink p-traps within a broader ecological context: they are habitats that repeatedly receive new microbial inputs while also retaining established residents. In such systems, community change may become increasingly shaped by what is already present, rather than by new input alone. Earlier colonists can occupy space, alter local conditions, or otherwise change the opportunities available to later arrivals^1, 2^. In microbial systems, this logic is closely related to the priority effects and the history-dependent nature of microbiome assembly. Our study did not directly test priority effects by experimentally varying the arrival order of defined colonists. However, it supports the narrower conclusion that p-trap communities became increasingly shaped by their prior state as succession progressed.

One useful way to interpret this system is through community coalescence^15^. In this view, each flushing event brought an incoming water-associated community into contact with the organisms already developing within the p-trap. In the early stage, when resident community was still forming, community composition remained more closely tied to recent input. Consistent with this, recent handwashing-water or tap-water input explained additional variation, and early-prioritized taxa included human- and water-associated organisms such as *Staphylococcus epidermidis, Staphylococcus hominis*, and *Cutibacterium acnes*. Later in succession, the community’s prior state became more informative than the most recent input, suggesting that incoming organisms were encountering an established resident community.The taxon-level attribution results supported this interpretation: early-phase taxa were more strongly associated with recent external input, whereas late-phase taxa were more strongly associated with the prior p-trap community state, consistent with increasing resident persistence. Thus, the dominant temporal axis reflects a shift in community organization, from early assembly shaped more strongly by recent input toward a later resident state shaped more strongly by prior community structure.

The short-timescale perturbation analysis provided an additional test of this interpretation. On the final day, flushing displaced communities from their immediate pre-flush positions in PCA space, showing that late-stage communities could still respond to disturbance. However, over the following hours, post-flush samples generally remained close to, or moved back toward, the late-stage resident centroid. Thus, flushing produced a short-term shift in community composition, but did not substantially move communities away from the late-stage state that had already formed. This supports the interpretation that late-stage p-trap communities were perturbable, but their short-term dynamics were constrained by the resident community that had already developed. Such behavior is consistent with community-level ecological memory-like dynamics^16, 17^.

This ecological framing also links our results to broader work on built-environment and premise-plumbing microbiomes. Recent reviews argue that built environments should be treated as distinct microbial ecological systems and also useful study systems because they are often more constrained than natural habitats and can be altered through design, materials, flow, human use, and cleaning^4^. Premise plumbing provides a particularly clear example: repeated wetting, biofilm-supporting surfaces, chemical heterogeneity, and strong spatial and temporal gradients create conditions in which resident microbial communities can develop^18^. Recent studies have shown that plumbing biofilm formation is dynamic and phase-structured^13^, and that sink p-trap communities can show long-term temporal dynamics and re-establishment after disturbance^14^. Our study extends those observations by showing how repeated input and local retention can interact over succession. Continued flushing does not necessarily keep p-trap communities closely tied to the most recent incoming water. Incoming communities may have their strongest influence during early assembly, before an established resident community increasingly shapes later community change.

Several limitations should be acknowledged. This was a controlled bench-scale system rather than a full occupied plumbing network, and that simplification was valuable precisely because it made assembly processes easier to interpret. The primary longitudinal analysis focused on p-trap water rather than continuous paired sampling of attached biofilm, so the inferred resident state may reflect a coupled water-phase and surface-associated system rather than water alone. However, our surface sampling at the end of the experiment suggests that very little biofilm was able to form during the limited time frame of the experiment. In addition, although the increasing explanatory value of each community’s previous state is consistent with growing historical dependence, our observational time-series design does not directly test whether arrival order causally altered community assembly. Future work that jointly tracks water and biofilm states, manipulates inoculum composition and arrival order, and tests stronger or repeated disturbances should help determine when resident persistence emerges, how general it is across built-environment habitats, and how strongly established communities constrain later colonization.

Overall, community development in newly prepared, bleach-treated p-traps was not simply a passive reflection of incoming water. Early in succession, recent external input explained a larger share of community variation, indicating that the habitat was still open to source-associated establishment. As succession progressed, however, p-trap communities became increasingly predictable from their own prior state and were only locally displaced by short-term flushing. These patterns indicate a shift from input-dominated early assembly toward resident persistence. More broadly, this work suggests that repeated input does not necessarily mean continued input control. Incoming communities may strongly shape early assembly, but their influence can decline as resident communities become established.

## Methods

### P-trap control experiment: apparatus setup, operation, and sampling

A bench-scale p-trap facility was constructed to allow longitudinal sampling of water from p-traps while simulating regular flushing from sink usage (Fig. S2). Each experimental unit was assembled using new components consisting of (i) an upper reservoir bucket connected to a faucet and equipped with a flow-control valve to deliver a steady inflow rate while minimizing splashing, (ii) a p-trap “reactor” (PVC p-trap) fitted with a removable sampling cap connected to an outlet tailpiece to facilitate aseptic sampling, and (iii) a lower collection bucket to capture wastewater. Prior to study initiation, all buckets and p-trap components were disinfected by overnight bleaching and thoroughly rinsed with tap water.

Three identical p-trap units were operated in parallel with regular flushing. Two units were flushed with handwashing water (experimental replicates), and one unit was flushed with tap water (control). Handwashing water was prepared by immersing both hands in 10 L of tap water and mixing the water by shaking the container; the mixed handwashing water was then split into two equal 5 L portions for the two replicate experimental units. Tap-water for flushing was collected directly from the same faucet that was used to prepare handwashing water. A flush consisted of 5 L of the assigned flush water per p-trap. Upper reservoir buckets were rinsed before preparation of flush water 10 L of tap water from the same faucet to minimize carryover and maintain comparable background conditions across units.

The experiment was conducted from May 7, 2024 to August 9, 2024. Initially (days 0 - 8), p-traps were flushed every 12 h by tap water. In the middle phase (days 9 - 73) p-traps were flushed twice per weekday, with no flushes on weekends. Two of the p-trap were flushed by handwashing water and one keep being flushed by tap water. In the last phase (days 74 - 94) flushing was reduced to once per weekday. P-trap water samples were collected using sterile syringes immediately prior to flushing. To homogenize the p-trap water before subsampling, the syringe was used to gently draw and expel liquid several times, after which 100 mL was collected using the syringe volume markings. Influent samples were collected in parallel: 500 mL of handwashing water (after mixing) and 500 mL of tap water were collected prior to flushing.

On the final sampling day, an intensive time-course was performed: p-trap water was collected immediately before flushing and at 0 h, 2 h, 6 h, and 12 h post-flush until the p-trap water was exhausted; following exhaustion, the inner surface of the p-traps was sampled with sterile swabs to collect any surface biofilms present.

### DNA extraction, library preparation and sequencing

All liquid samples were filtered through 0.2 μm membrane filters. Filters and swab heads (from final day surface samples) were aseptically cut into small pieces using disposable sterile scalpels prior to DNA extraction. Genomic DNA was extracted using the DNeasy PowerSoil Pro kit (QIAGEN) following the manufacturer’s instructions, and DNA concentration was quantified using a Qubit fluorometer with dsDNA Quantification HS Assay Kits. Our lower limit of detection for DNA was 0.05 ng*/*μL because only 2 μL per sample was used for DNA quantification in order to preserve limited sample.

Both amplicon and shotgun metagenomic sequencing strategies were used for this study. All samples were profiled by PacBio full-length 16S rRNA gene amplicon sequencing. For 16S rRNA sequencing, extracted DNA was submitted to the Duke Microbiome Core, where full-length 16S rRNA gene PCR amplification, library preparation, and PacBio sequencing were performed, as follows, using the PacBio Kinnex 16S kit (PacBio). Total DNA extracted from each sample was quantified via Qubit and normalized to 0.2 ng*/*μL for each sample. 1 ng of DNA was used as template for PCR amplification of the full-length 16S rRNA gene, using Phusion Plus PCR Master Mix (ThermoScientific) with the 27F-1492R universal primer set (5’-AGRGTTYGATYMTGGCTCAG-3’ and 5-RGYTACCTTGTTACGACTT-3’, respectively) containing unique barcodes and Kinnex adaptors at a final concentration of 0.3 μM. Reaction mixes were denatured for 30 secs at 98 °C prior to 25 cycles for high concentration samples and 35 cycles for low concentration samples of denaturation at 98 °C for 10 seconds, annealing at 57 °C for 20 seconds, and extension at 72 °C for 75 seconds. After the last cycle, a final extension at 72 °C for 5 minutes was performed. Completed PCR reactions were visualized on an E-Gel (Invitrogen) to ensure that amplicon size was correct (1500bp), and that each sample amplified appropriately. Amplicon libraries were subsequently pooled without cleanup at varying volumes based on gel band intensity (1, 2.5, 5, 10, or 20 μL per reaction). Library pool was cleaned and concentrated using 1.1x volume of SMRTbell® cleanup beads (PacBio) and eluted in 200 μL or 100 μL prior to Kinnex PCR for concatenation and circularization. Kinnex PCR was performed as outlined in the PacBio Kinnex 16S kit’s published protocol with no modifications (PacBio). Size selected and cleaned libraries were loaded onto a PacBio SMRT® Cell and sequenced on the Revio system (PacBio) in the Duke Sequencing and Genomic Technologies Shared Resource.

To monitor background contamination and amplification performance, multiple blank and control samples were included during sample processing and sequencing. Extraction blanks were generated by processing molecular-grade water through DNA extraction in place of a biological sample. The Duke Microbiome Core also included PCR negative and positive controls during amplification. All blanks and PCR negative controls were used subsequently for ASV-level contaminant identification.

Shotgun metagenomic sequencing was done for a subset of the samples (samples collected on Mondays from June 17 to July 22) that had biomass high enough to yield a sufficient amount of DNA. Shotgun metagenomic libraries were prepared using the ligation-based Native Barcoding Kit (SQK-NBD114; Oxford Nanopore Technologies) and sequenced on a PromethION 2 Solo (Oxford Nanopore Technologies) using R10.4.1 PromethION flow cells.

### Processing and curation of PacBio full-length 16S datasets

Raw PacBio HiFi full-length 16S reads were processed into ASV tables using the Pacific Biosciences HiFi 16S workflow^1^, which implements DADA2-based denoising^19^. We used a modified version of the workflow in which the maximum expected error threshold was set to 3. Low-abundance ASVs (ASVs with 20 or less counts combined across the batch) were removed before downstream analysis. Count tables were converted to relative-abundance matrices for ecological summaries and to centered log-ratio (CLR) coordinates for multivariate analyses. For CLR transformation, zeros were replaced with a small offset defined as half of the smallest positive value observed in the matrix. CLR transformation was used because amplicon data are compositional and are therefore more appropriately analyzed in ratio space than as unconstrained counts or proportions^20–22^.

### Blank-aware decontamination and confidence filtering

Because contamination is a central concern in low-biomass microbiome studies, the PacBio full-length 16S dataset was subjected to explicit blank-aware decontamination. Contaminant ASVs were identified using the prevalence-based approach implemented in decontam^23^, and a decontam threshold of 0.5 was used for downstream filtering.

After ASV-level decontamination, sample-level confidence was assessed using post-cleaning read depth, the fraction of reads retained after decontamination, and residual read counts after decontamination. Samples were classified as low-confidence if, after cleaning, read depth was less than the median read depth of PCR negative controls, if <25% of reads were retained after decontamination, or if the cleaned sample contained <1,000 reads. All downstream analyses were then performed on the decontamination and high confidence subset.

### Robustness of community profiles across PCR cycle conditions

To assess robustness across amplification conditions, we analyzed the subset of biological samples that had been sequenced using both the 25-cycle and 35-cycle workflows. ASV tables were aligned by shared feature identity, missing feature values were set to zero, and the combined matrices were transformed to CLR coordinates. Matched 25-cycle and 35-cycle samples were then compared using Spearman correlation on ASV-level CLR profiles.

We additionally assessed concordance in temporal ASV behavior across PCR conditions. The 200 most abundant ASVs in the merged dataset were selected based on mean relative abundance, and for each ASV, temporal association was evaluated separately within the 25-cycle and 35-cycle datasets using linear models with CLR-transformed abundance as the response, day of year as a continuous predictor, and p-trap identity as a categorical fixed effect to account for p-trap-specific baseline differences. P values for the day-of-year term were adjusted across ASVs using the Benjamini–Hochberg procedure. Signed time-trend score was defined as sign(estimate) × -log10(p). Concordance in both effect direction and statistical support between the 25-cycle and 35-cycle analyses was used as a complementary measure of cross-batch reproducibility.

### Compositional ordination of p-trap communities

Community composition was analyzed by principal component analysis (PCA) on CLR-transformed ASV tables. PCA on CLR coordinates was used to obtain an interpretable Euclidean representation of compositional community variation while avoiding distortions associated with direct analysis of raw proportions^20, 21^. We first performed a global PCA using combined datasets to visualize the major structure across sample types and PCR conditions. This analysis was used to compare overall community organization and to examine matched 25-cycle and 35-cycle samples in a shared ordination space. We then performed separate PCA analyses restricted to the p-trap subset within each dataset to resolve longitudinal structure within the experimental p-traps without the broader between-sample-type variation dominating the ordination. In all cases, the first two principal components were used for visualization.

### Cross-platform validation using overlapping metagenomic and 16S datasets

To assess whether the temporal patterns identified in the 16S dataset were independently recapitulated by shotgun metagenomic profiling, we analyzed an overlapping set of weekly samples from handwashing p-traps collected from mid-June to late July. Metagenomic samples were matched to 16S samples by date and p-trap identity, and comparisons were restricted to the matched subset. Taxonomic profiles from both platforms were aggregated at the genus level, and only genera detected in both datasets were retained for downstream cross-platform analysis.

To examine whether both datasets captured a common temporal signal, shared-genus abundance tables were transformed to centered log-ratio (CLR) coordinates after replacing zeros with a pseudocount equal to half of the minimum positive abundance in the corresponding matrix, and principal component analysis (PCA) was performed separately for the 16S and metagenomic datasets. The sign of the first principal component (PC1) was oriented such that increasing PC1 corresponded to later sampling dates. Association between PC1 and time was then evaluated using Spearman correlation.

To identify taxa associated with the temporal axis in each platform, genus-level CLR abundances were correlated with platform-specific PC1 scores using Spearman correlation, yielding one genus-specific association coefficient per platform. Cross-platform agreement in temporally associated genera was then assessed across the shared genus set by comparing these genus-specific PC1 association coefficients between the 16S and metagenomic datasets. In addition to correlation of effect sizes across genera, sign concordance was calculated as the proportion of shared genera for which the PC1 association had the same direction in both platforms, that is, positive in both or negative in both. This analysis was used to determine whether the dominant temporal structure and its principal genus-level drivers were consistently recovered across sequencing platforms.

### Short-term projection analysis of final-day post-flush dynamics

To assess short-term response of established p-trap communities to flushing, we projected the intensive final-day post-flush samples into the fixed PCA space defined by the original 25-cycle p-trap ordination. Although some final-day timed samples were also present in the 35-cycle dataset, those data were incomplete across p-traps and post-flush timepoints; therefore, this analysis was performed on the 25-cycle dataset only. Pre-flush samples collected immediately before flushing were used as within-p-trap reference points, and a late-stage resident centroid was calculated for each p-trap from the three most recent pre–final-day samples in the original PCA scores. Euclidean distances from each projected timed sample to the pre-flush sample and late-stage centroid were then calculated in PCA space to assess short-term displacement and partial recovery toward established resident community structure.

### Temporal modeling of community succession along major compositional axes

To summarize temporal community dynamics beyond static ordination, we applied a custom PCA-axis longitudinal framework to the p-trap datasets. Briefly, PCA was performed on CLR-transformed ASV tables, and downstream longitudinal analyses were restricted to the first two principal components, which captured the dominant axes of community variation.

Temporal structure along the leading PCA axes was modeled using generalized additive models (GAMs) with the time smoother specified using *k* = 6, allowing nonlinear changes in community state to be estimated flexibly over time^24^. These analyses were used to summarize treatment-level temporal trajectories and overall axis-shift patterns, thereby assessing whether the dominant axes of compositional change were temporally structured and reproducibly associated with succession.

### Nested comparison of external-input versus lagged within-p-trap explanatory gain

To assess whether the dominant ecological control on p-trap community composition changed across succession, we compared the explanatory gain provided by recent external input versus lagged within-p-trap state using a nested-model framework in CLR space. Because the aim of this analysis was to determine whether a focal community state was better explained by recent incoming water or by the p-trap’s own immediately preceding state, rather than to estimate literal source fractions, we quantified model improvement relative to a blank/background baseline rather than applying conventional source-tracking proportions.

Taxa were first filtered to retain reproducibly observed features (prevalence ≥ 5% and mean relative abundance ≥ 1 × 10^*−*4^). Relative-abundance profiles were then transformed to centered log-ratio (CLR) coordinates. For each focal p-trap sample, candidate external samples were restricted to those collected before the focal sample and within a predefined look-back window, which was 1 day in the primary analysis. In treated p-traps, recent handwashing-water samples were used to define the external-input profile, whereas in the control p-trap recent tap-water samples were used instead. The immediately preceding sample from the same p-trap was used to represent lagged within-p-trap state, and a blank-derived profile was included to represent background low-biomass structure. External profiles were summarized from eligible prior samples as a CLR-space summary profile. Sensitivity analyses repeated the same nested-model procedure using alternative external-profile definitions, including a 3-day summary profile and a 3-day time-decayed profile in which more recent eligible external samples received greater weight.

For each focal sample, we fit four predefined nested models in CLR space: a blank-only model (*M*_blank_), an external-plus-blank model (*M*_ext_), a lag-plus-blank model (*M*_lag_), and a full model including blank, external input, and lagged within-p-trap state (*M*_full_). Ridge regularization was used to stabilize estimation when external and lagged source profiles were partially correlated. Model fit was summarized by CLR-space *R*^2^.

The contrasts used for the main phase comparison analysis were the incremental gains in explanatory power relative to the blank baseline:

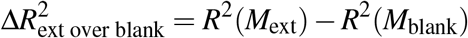

and

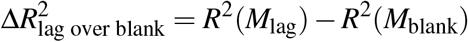

Positive values therefore indicate that adding the corresponding term improved explanation of the focal community beyond the blank/background baseline. These per-sample Δ*R*^2^ values were compiled in the model-comparison table and then visualized by temporal phase to compare whether recent external input or lagged within-p-trap state contributed greater explanatory gain at different stages of succession. Temporal phases were defined a priori by days since the start of the p-trap time series as early (days 0-14), mid (days 15-35), and late (days *>* 35). We explored a data-driven alternative based on segmented regression of sample-level support for external input versus lagged within-p-trap state, but the estimated breakpoints varied across PCR-cycle subsets and p-trap groups and were therefore not sufficiently stable for a common phase definition. In the 25-cycle subset, bootstrap confidence intervals were especially broad, owing largely to sparse early external-support signal in p-trap 3 rather than to a clear transition in community source attribution. Because these inferred breakpoints were sensitive to sample composition and did not yield a robust, comparable partition across datasets, we retained the fixed day-based phase definitions for downstream analyses. Figure 5 summarizes the distribution of sample-level Δ*R*^2^ values within each phase, with each point representing one focal p-trap sample. Corresponding sensitivity plots based on alternative external-profile definitions were used to confirm that the phase-dependent contrast between external-input gain and lag-based gain was qualitatively robust to the definition of recent external input.

For phase-level statistical comparisons shown above the boxplots, we performed early-versus-late comparisons within each plotted panel. Specifically, sample-level Δ*R*^2^ values were compared between the early phase (days 0-14) and the late phase (days *>* 35) using a two-sided Wilcoxon rank-sum test. The same procedure was applied separately for each model contrast, including the external-input gain and lagged within-p-trap gain contrasts, and was repeated independently for each sensitivity-analysis panel.

### Taxonomic attribution of early colonization and late persistence

To identify taxa associated with early external colonization versus later within-p-trap persistence, we extended the nested-model framework described above to the feature level and then summarized the resulting attribution statistics by taxon. Using the same filtered feature set, source definitions, and phase assignments as in the sample-level analysis, we extracted feature-level fitted values and squared residual errors under the four nested models (blank-only, external-plus-blank, lag-plus-blank, and full external-plus-lag-plus-blank) for each focal p-trap sample.

For each feature *f* in each focal sample *i*, we defined the squared residual error under model *M* as

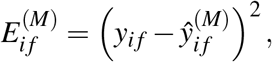

where *y*_*i f*_ is the observed CLR abundance and 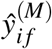 is the fitted value under the corresponding nested source model. Feature-level attribution was then quantified as the reduction in squared residual error achieved by a more complex model relative to a simpler nested alternative. Specifically, we calculated

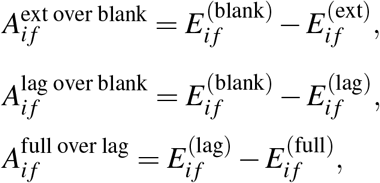

and

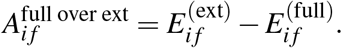

Positive values therefore indicate that inclusion of the additional source term improved explanation of that feature’s CLR abundance in the focal sample.

These feature-level attribution values were then aggregated by taxon within phase across all contributing feature-sample observations. Early external-associated taxa were ranked using the summed external-over-blank attribution in the early phase, whereas late history-associated taxa were ranked using the summed lag-over-blank attribution in the late phase. For visualization, we plotted the top-ranked early-colonization and late-persistence taxa from the 25-cycle dataset and from the 35-cycle dataset separately.

## Data availability

The sequence data have been deposited with links to BioProject accession number XXXXX in the NCBI BioProject database.

## Code availability

All analyses are fully reproducible using the code available in the accompanying GitHub repository (https://doi.org/XXXXXXXXXXX), the publicly available Apptainer images indicated in the code, and the publicly available data available as discussed above.

## Additional information Acknowledgements

This work was supported by the Engineering Research Centers Program of the National Science Foundation under NSF Cooperative Agreement No. EEC-2133504. Any opinions, findings and conclusions or recommendations expressed in this material are those of the author(s) and do not necessarily reflect those of the National Science Foundation.

We thank the Duke University School of Medicine for the use of the Microbiome Core Facility, which provided Microbiome 16S rRNA library service, and for the use of the Sequencing and Genomics Core Facility for PacBio full-length 16S rRNA sequencing service. Purchase of the PacBio Revio was funded by the NIH (1S10OD034222-01).

## Footnotes

1 https://github.com/pacificbiosciences/HiFi-16S-workflow

