## Supplementary Information for "From external-input sensitivity to resident persistence: community assembly in a sink p-trap model"

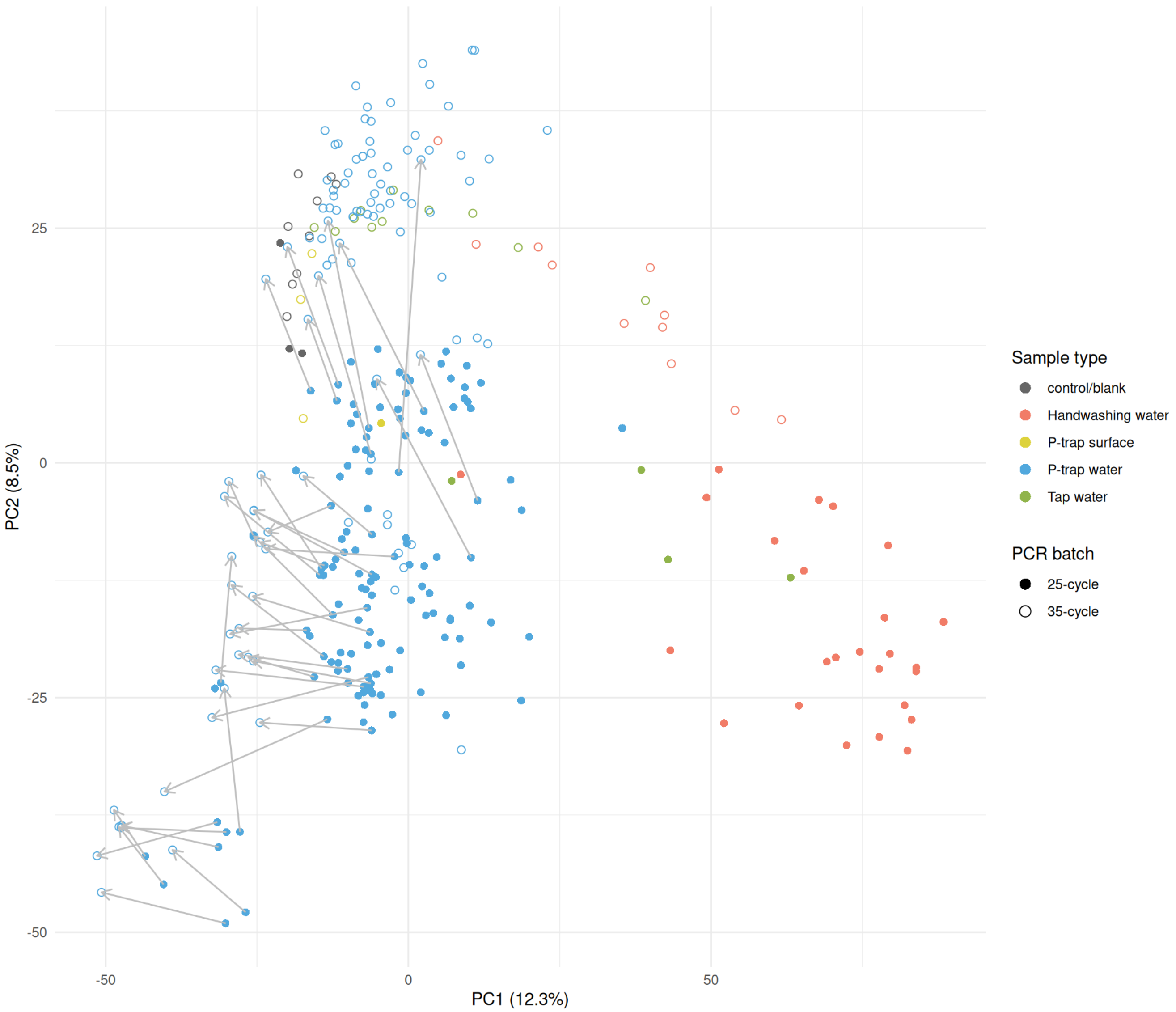


**Figure S1.** Global PCA shows preservation of sample type level community structure across PCR-cycle conditions. Colors indicate sample type: control/blank (gray), handwashing water (salmon), P-trap surface (yellow), P-trap water (blue), and tap water (green). Point style indicates PCR batch: filled circles represent 25-cycle samples, and open circles represent 35-cycle samples. Gray lines connect the matched samples selected for sequencing in both the 25-cycle and 35-cycle batches.

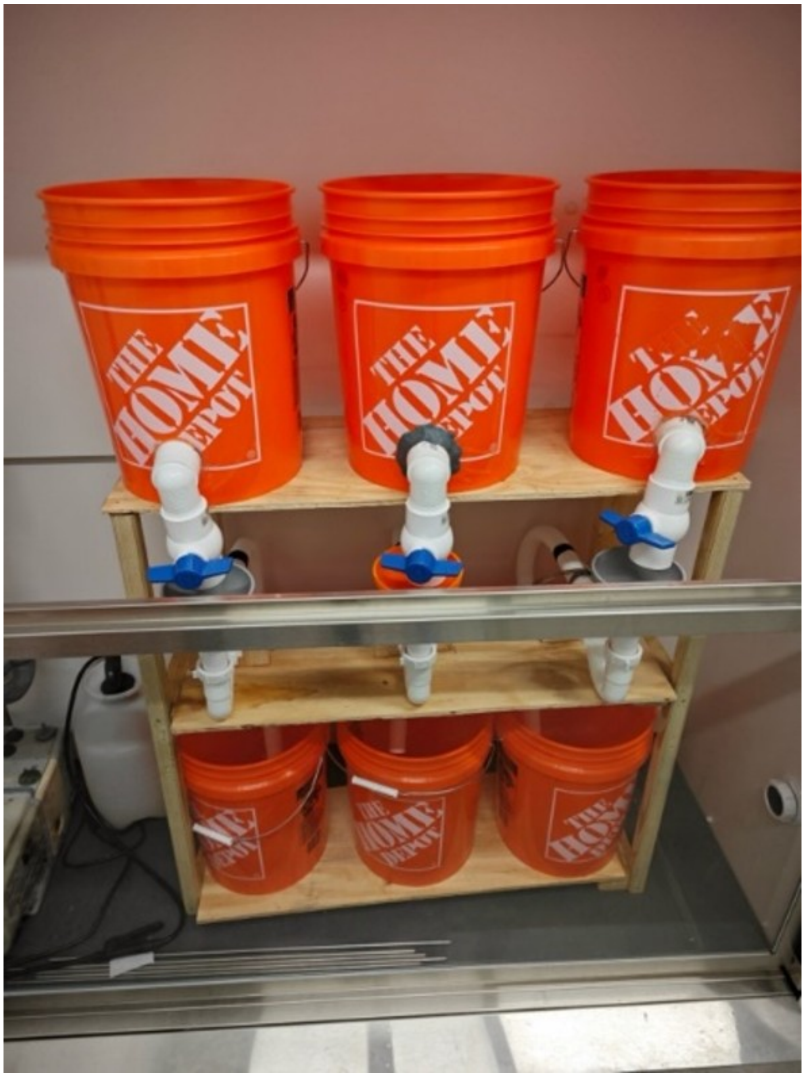


**Figure S2.** Bench-scale p-trap facility used for the p-trap experiment.

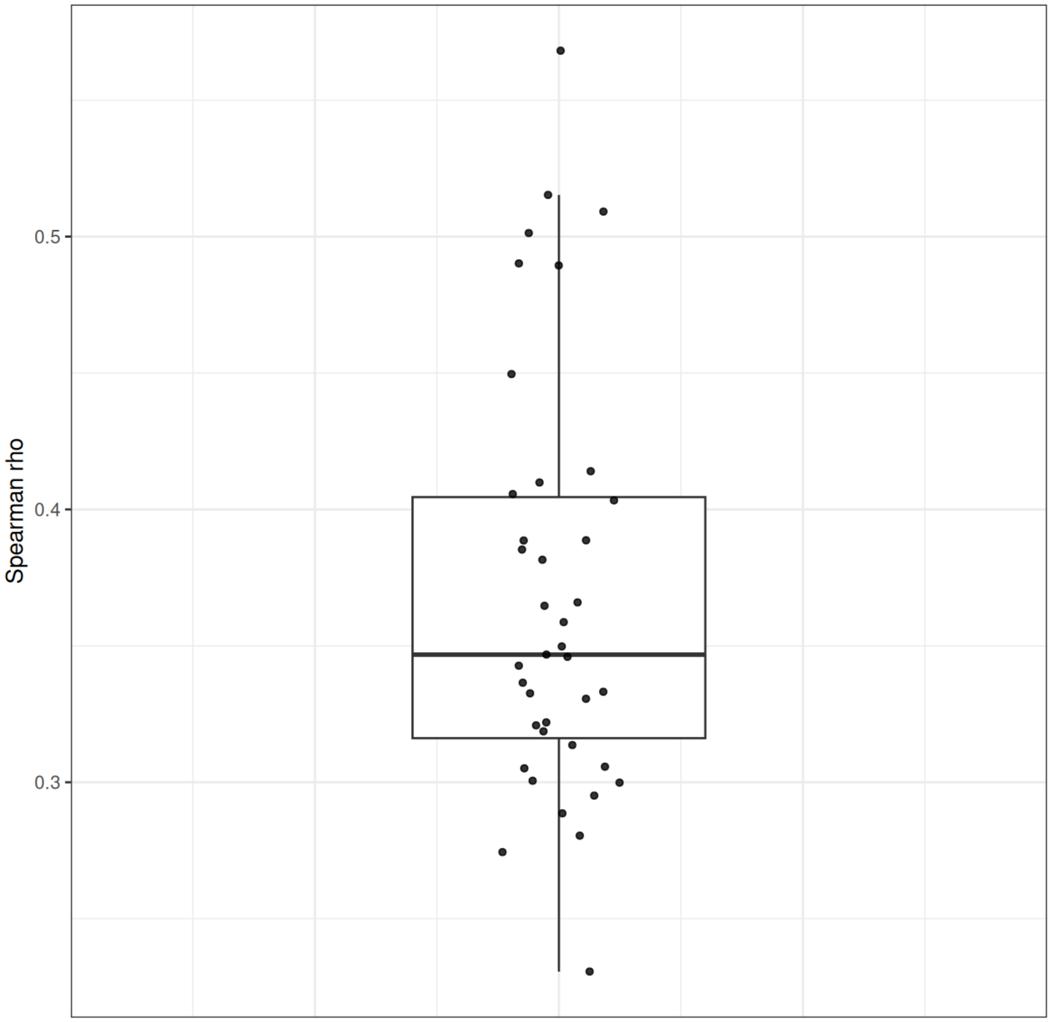


**Figure S3.** Matched samples showed moderate but consistent profile-level concordance across PCR-cycle conditions. For each matched 25-cycle and 35-cycle sample pair, concordance was quantified as the Spearman correlation of ASV-level CLR abundance profiles. Points represent individual matched pairs, and the boxplot summarizes the overall distribution of pairwise correlations.

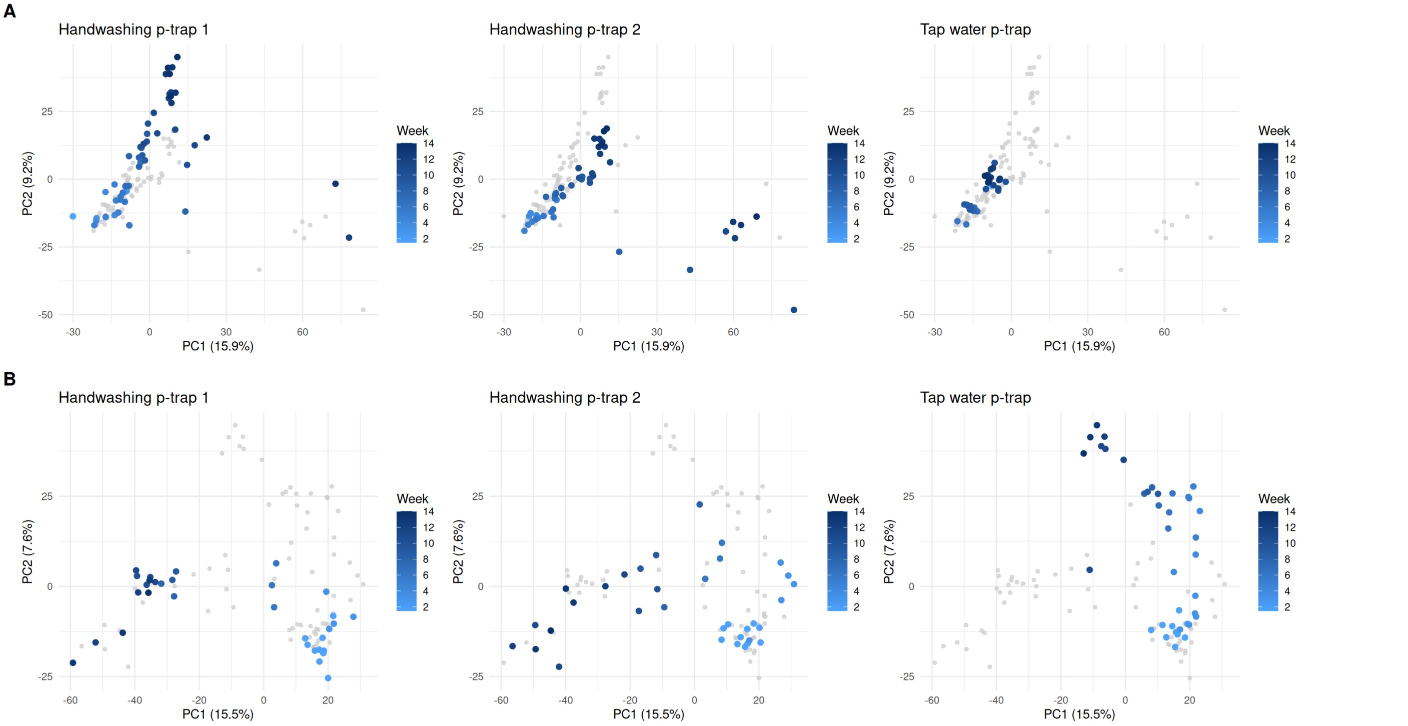


**Figure S4.** Directional successional trajectories of p-trap communities in PCA space across PCR-cycle datasets. (A) 25-cycle dataset. (B) 35-cycle dataset. Points are colored by sampling week and faceted by p-trap.

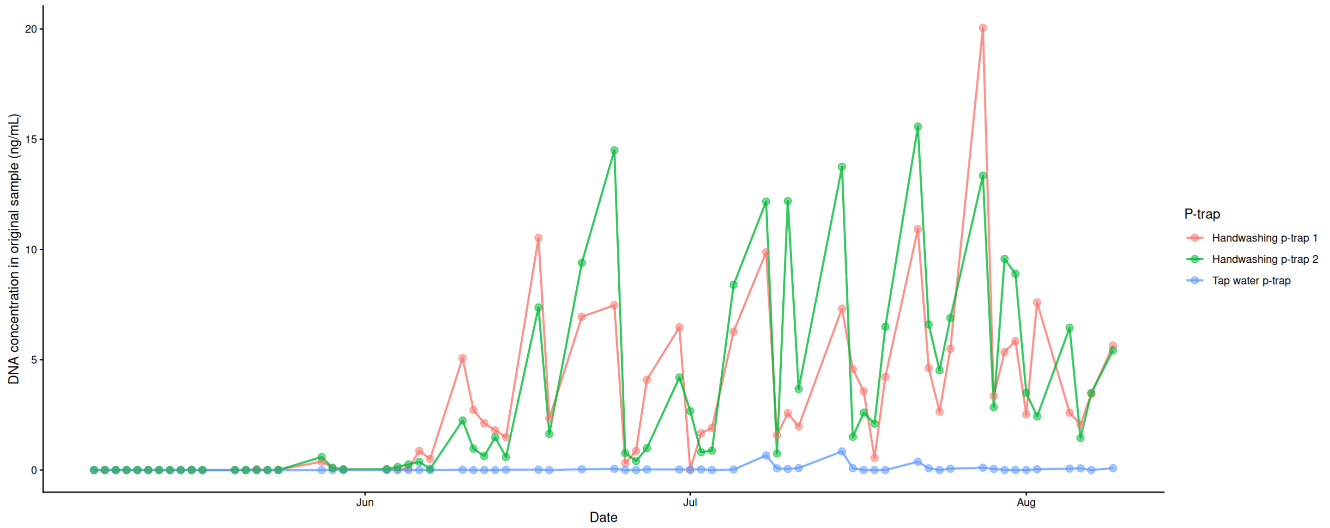


**Figure S5.** The estimated DNA concentration of p-trap water samples along time.

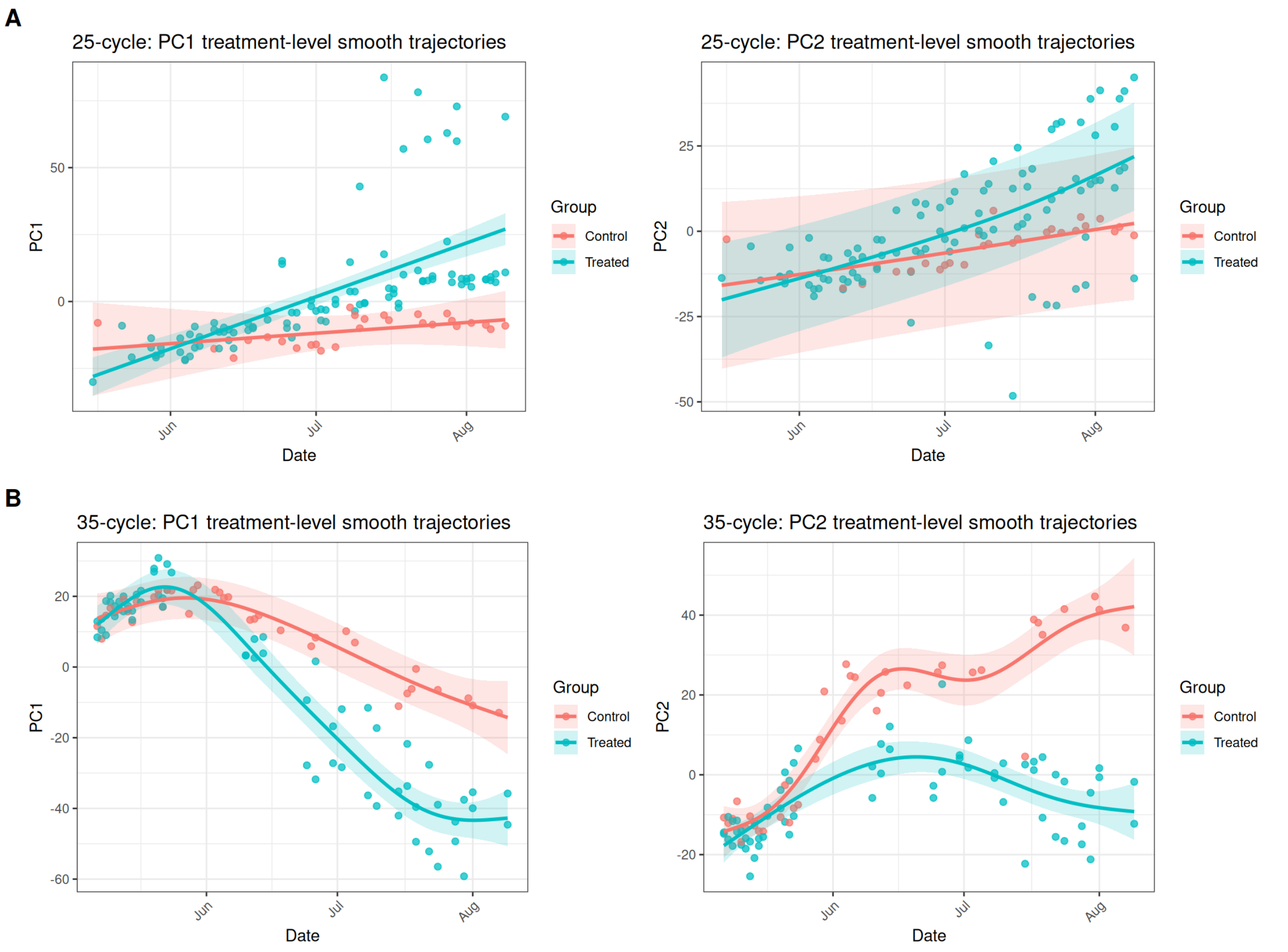


**Figure S6.** Treatment-level temporal trajectories along the major ordination axes. Points show sample scores, and lines with shaded bands show GAM-fitted trajectories ±95% confidence intervals for treated and control p-traps.

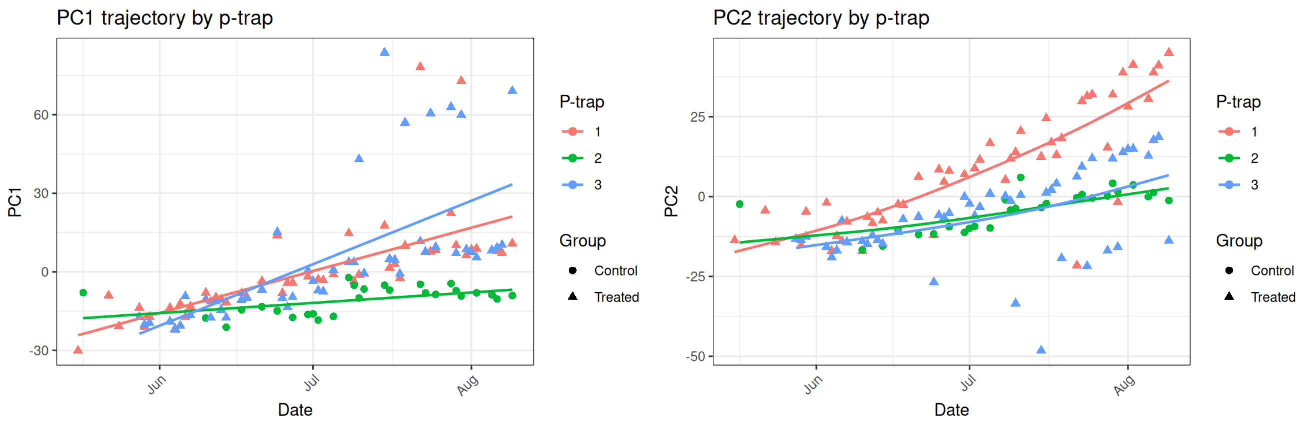


**Figure S7.** Temporal trajectories along the major ordination axes by different p-traps in 25 cycle dataset. P-trap 1 and 3 are flushed by handwashing water and p-trap 2 is flushed by tap water.

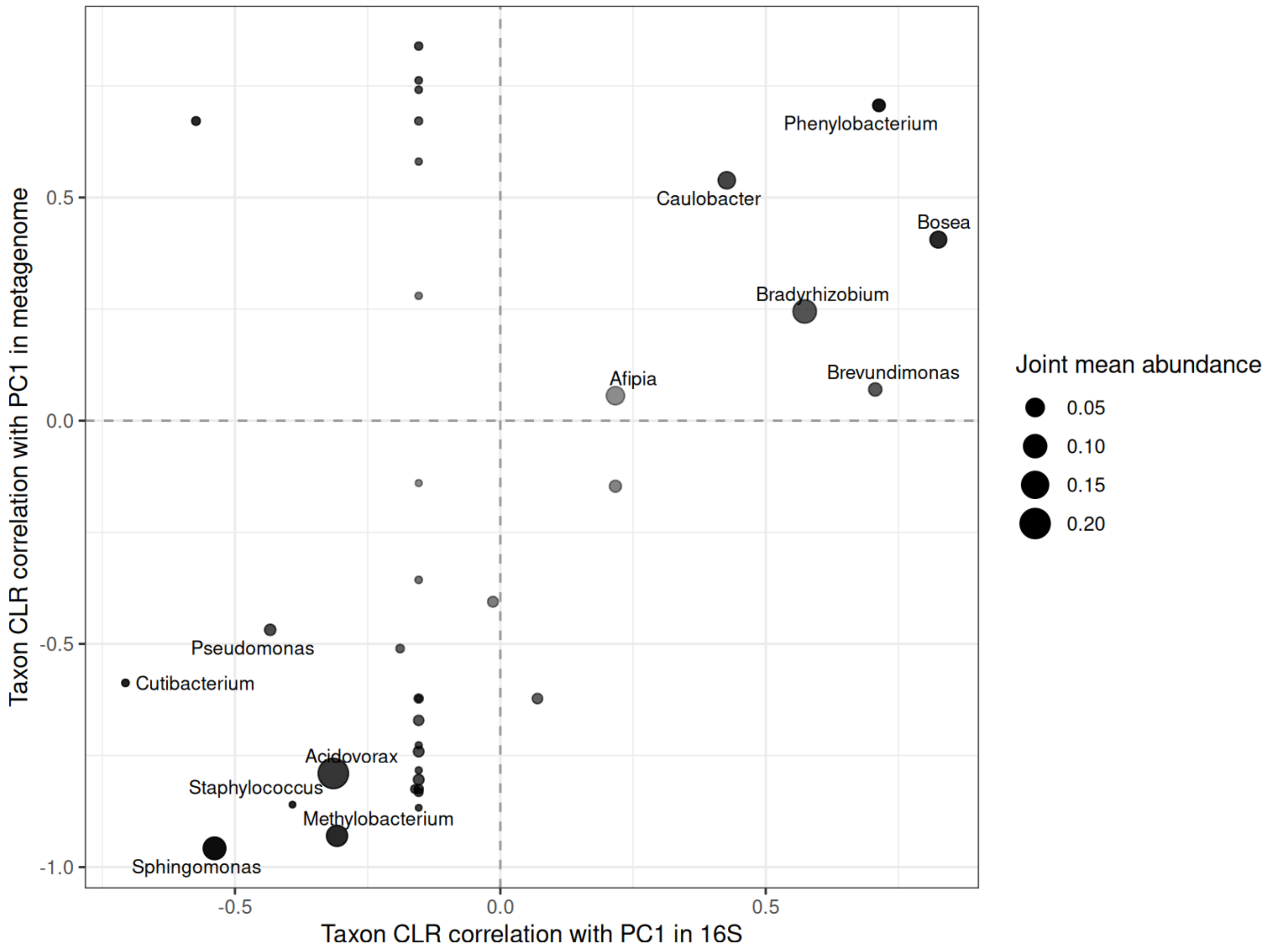


**Figure S8.** Shared genera showed concordant association with the dominant temporal axis across sequencing modalities. Each point represents a shared genus; axes show the correlation between genus-level CLR abundance and PC1 in the 16S and metagenomic datasets, respectively. Point size indicates joint mean abundance.

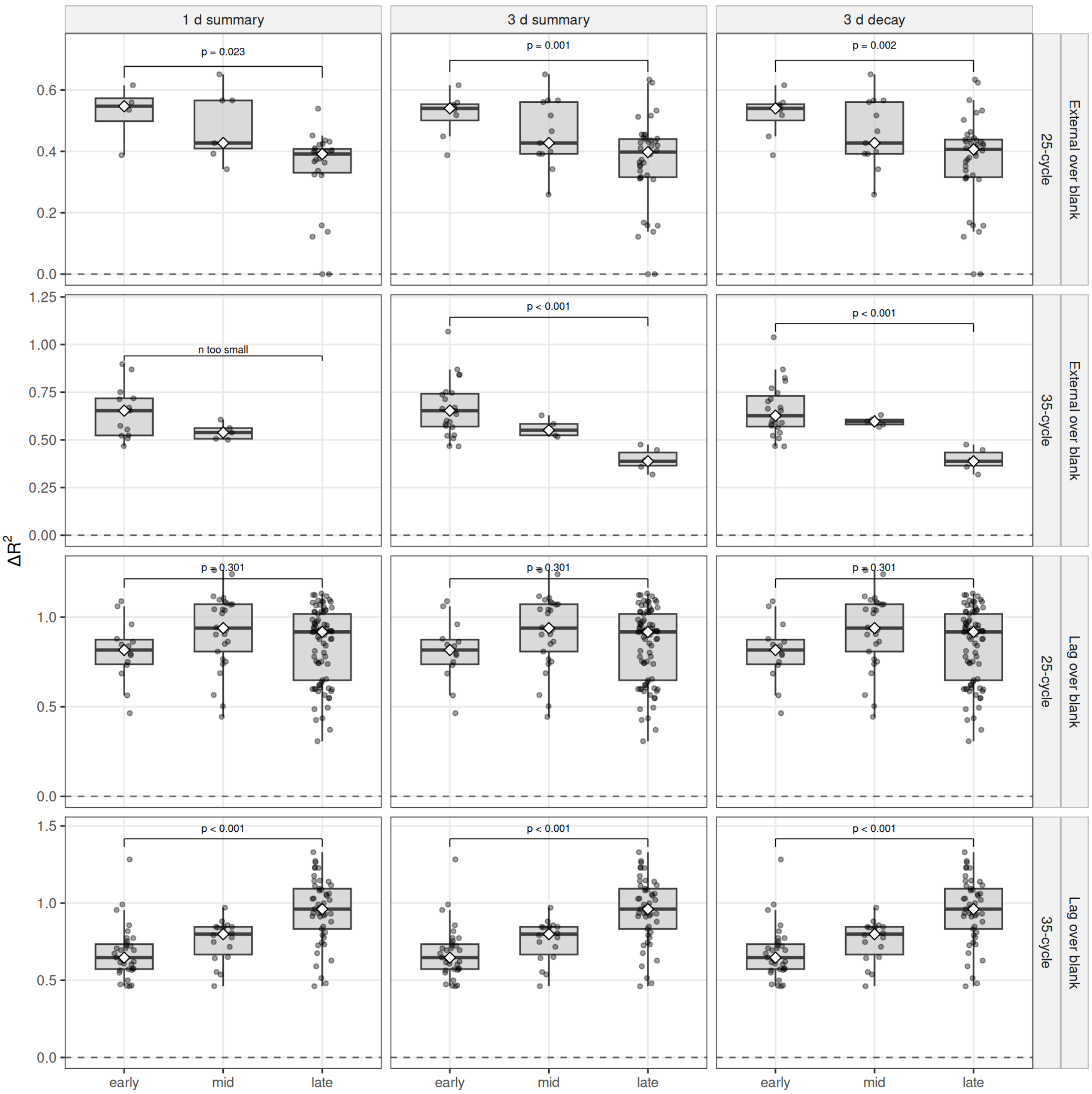


**Figure S9.** Phase-specific nested-model $\Delta R^2$ patterns were robust to alternative definitions of recent external input.For each focal sample, explanatory gain was quantified relative to the blank-only baseline as $\Delta R^2_{\text{ext over blank}}$ or $\Delta R^2_{\text{lag over blank}}$. Columns show alternative external-source definitions used in the sensitivity analysis (1-day summary, 3-day summary, and 3-day decayed profile), and rows show the 25-cycle and 35-cycle datasets. Across specifications, recent external input showed its strongest explanatory gain in early samples and approached zero in later phases, whereas lagged within-p-trap state showed positive explanatory gain throughout succession and became strongest in late samples. Although effect sizes varied across datasets and external-source definitions, the overall phase-dependent contrast was preserved, supporting a shift from early external-input sensitivity toward stronger late-stage within-p-trap persistence.

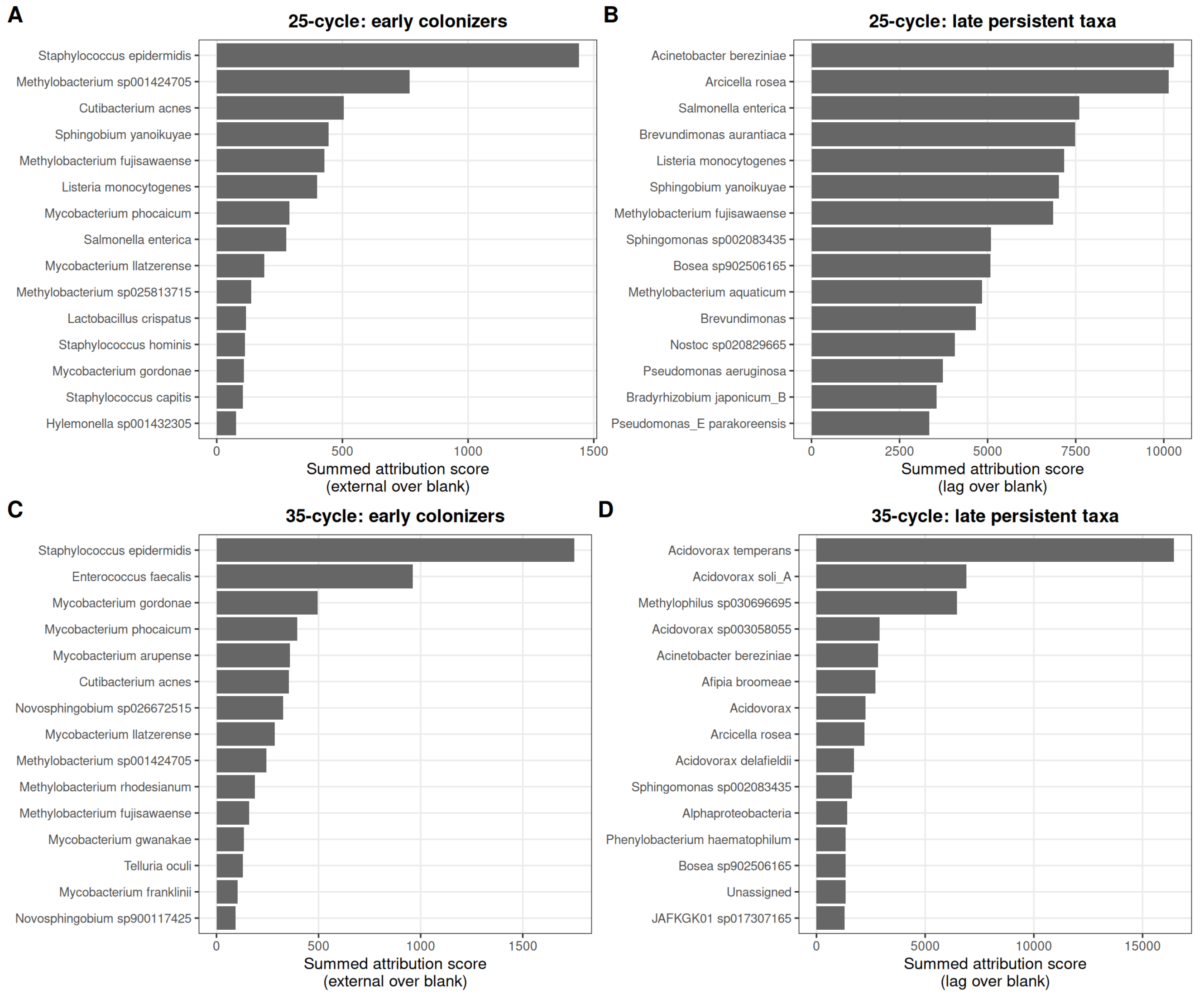


**Figure S10.** Taxon-level attribution highlights early colonizers and late persistent taxa in handwashing p-traps in 35-cycle dataset.Taxa are ranked by summed feature-level attribution scores within each dataset and phase.

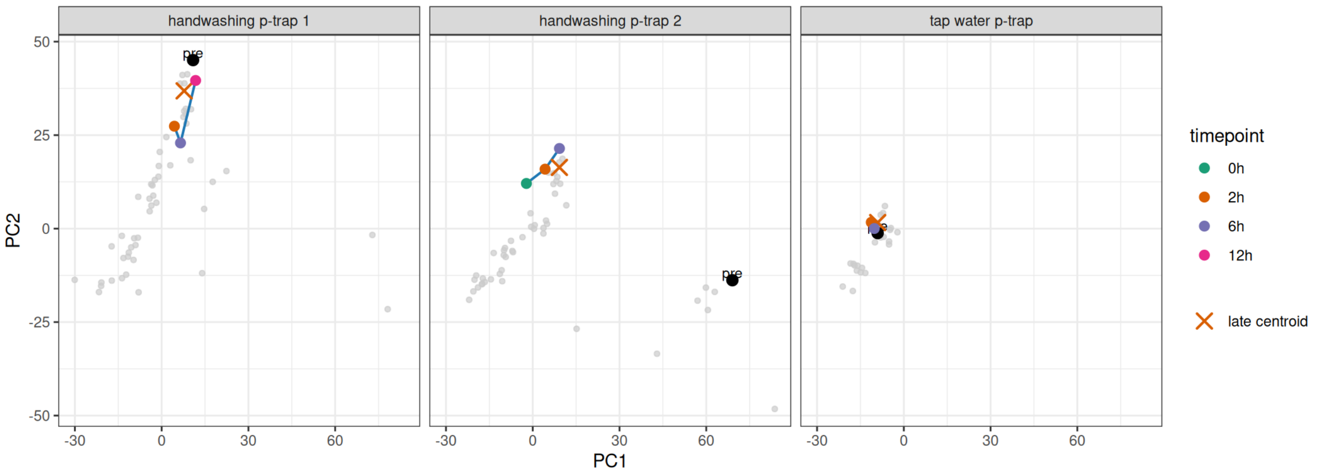


**Figure S11.** Final-day flush produced short-term displacement around late-stage resident community states. Timed post-flush samples collected on the final day were projected into the fixed PCA space defined by the original 25-cycle p-trap-only ordination, without refitting the PCA. Black points indicate the corresponding original pre-flush samples already present in the ordination, whereas colored points indicate projected post-flush samples collected at 0, 2, 6, and 12 h after flushing. Black circle shows pre-flush sample, orange "X" shows centroid of the three most recent longitudinal samples collected before the final-day recovery experiment. Although handwashing p-trap 2 starts from a more separated pre-flush position in PCA space, it still returns toward the late-stage centroid after flushing.

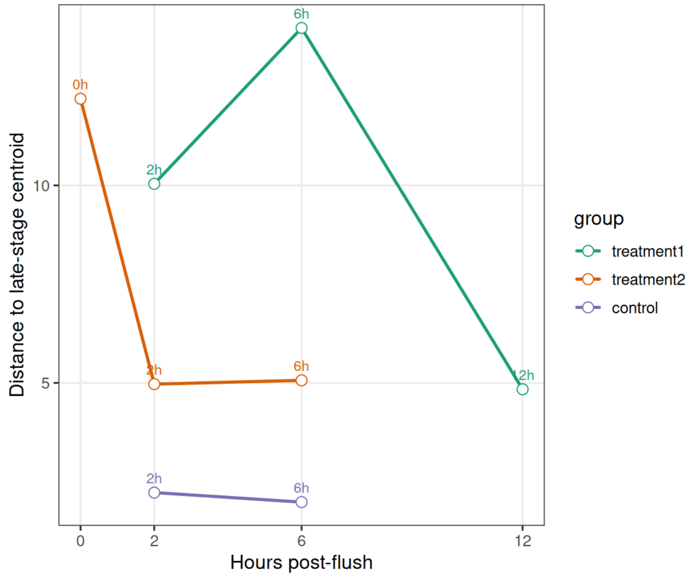


**Figure S12.** Distance of final-day timed samples to the late-stage resident centroid after flushing for 25 cycle data
